# Massively parallel identification of zipcodes in primary cortical neurons

**DOI:** 10.1101/2021.10.21.465275

**Authors:** Nicolai von Kügelgen, Samantha Mendonsa, Sayaka Dantsuji, Maya Ron, Marieluise Kirchner, Nadja Zerna, Lucija Bujanic, Philipp Mertins, Igor Ulitsky, Marina Chekulaeva

## Abstract

Cells adopt highly polarized shapes and form distinct subcellular compartments largely due to the localization of many mRNAs to specific areas, where they are translated into proteins with local functions. This mRNA localization is mediated by specific cis-regulatory elements in mRNAs, commonly called “zipcodes.” Their recognition by RNA-binding proteins (RBPs) leads to the integration of the mRNAs into macromolecular complexes and their localization. While there are hundreds of localized mRNAs, only a few zipcodes have been characterized. Here, we describe a novel <u>n</u>euronal <u>z</u>ipcode identification <u>p</u>rotocol (N-zip) that can identify zipcodes across hundreds of 3’UTRs. This approach combines a method of separating the principal subcellular compartments of neurons – cell bodies and neurites - with a massively parallel reporter assay. Our analysis identifies the *let-7* binding site and (AU)n motif as *de novo* zipcodes in mouse primary cortical neurons and suggests a strategy for detecting many more.

## INTRODUCTION

Many cell types deliver mRNAs to specific subcellular locations as a key mechanism to produce local pools of proteins that have a range of functions. This process supports, for example, the establishment of the body axis, cell growth and migration, and processes of learning and memory (reviewed in Holt et al., 2019; Martin and Ephrussi, 2009). It is particularly important in highly polarized cells such as neurons, whose functions depend on specific patterns of mRNA localization to cell bodies (soma) and extensions (neurites). The localization is thought to be primarily mediated by cis-regulatory elements (“zipcodes”) that are usually found in the 3’UTRs of the mRNAs (Kislauskis and Singer, 1992). Zipcodes are bound by specific RNA-binding proteins (RBPs) that link their targets to transport machinery or regulators of mRNA stability and direct their localization to the sites of function. Specific zipcodes and bound RBPs are often conserved between diverse cell types and species, supporting the idea that mRNA localization is an ancient and fundamental feature of development. As an example, the localization of beta-actin mRNA has important functions in ascidian eggs, chicken fibroblasts, and mouse neurons (Bassell et al., 1998; Jeffery et al., 1983; Kislauskis et al., 1994; Micheva et al., 1998).

High-throughput analyses have demonstrated specific localization patterns for hundreds to thousands of mRNAs in diverse organisms and cell types (Briese et al., 2016; Cajigas et al., 2012; Ciolli Mattioli et al., 2019; Gumy et al., 2011; Jambor et al., 2015; Lecuyer et al., 2007; Maciel et al., 2018; Poon et al., 2006; Shigeoka et al., 2016; Taliaferro et al., 2016; Tushev et al., 2018; Zappulo et al., 2017; Zhong et al., 2006; Zivraj et al., 2010). Presumably many of these events rely on a similar mechanism, but to date only a few zipcodes have been characterized. Here, we report the development of a method to systematically map and characterize neuronal zipcodes transcriptome-wide. Our approach combines a massively parallel reporter assay with the isolation of neuronal subcellular compartments – soma and neurites. Specific results include the identification of the *let-7* binding site and (AU)n motif as *de novo* zipcodes with functions in mouse primary cortical neurons.

## RESULTS

### Establishment of the neuronal zipcode identification protocol (N-zip)

To perform an unbiased transcriptome-wide analysis of zipcodes, we developed the <u>n</u>euronal <u>z</u>ipcode identification <u>p</u>rotocol (N-zip). This combines a <u>m</u>assively <u>p</u>arallel reporter <u>a</u>ssay (MRPA, Lubelsky and Ulitsky, 2018) with a neurite/soma fractionation scheme established previously in our lab (Ciolli Mattioli et al., 2019; Ludwik et al., 2019; Zappulo et al., 2017) (**Figure 1A**). In short, this method involves the following steps: (1) generating a set of fragments tiled across 3’UTRs of neuritically localized transcripts; (2) cloning these fragments into a vector library between adaptor sequences and delivery into neurons; (3) separating the soma and neurites of neurons; (4) preparing RNA-seq libraries from RNA fragments flanked by adapters; (5) identifying fragments that are sufficient for the localization of RNAs to neurites – indicating the potential presence of zipcodes; (6) carrying out an extensive mutagenesis of fragments that specifically localize to neurites, then repeating steps 2-5 to map the specific sequences that serve as zipcodes.

**Figure 1.**
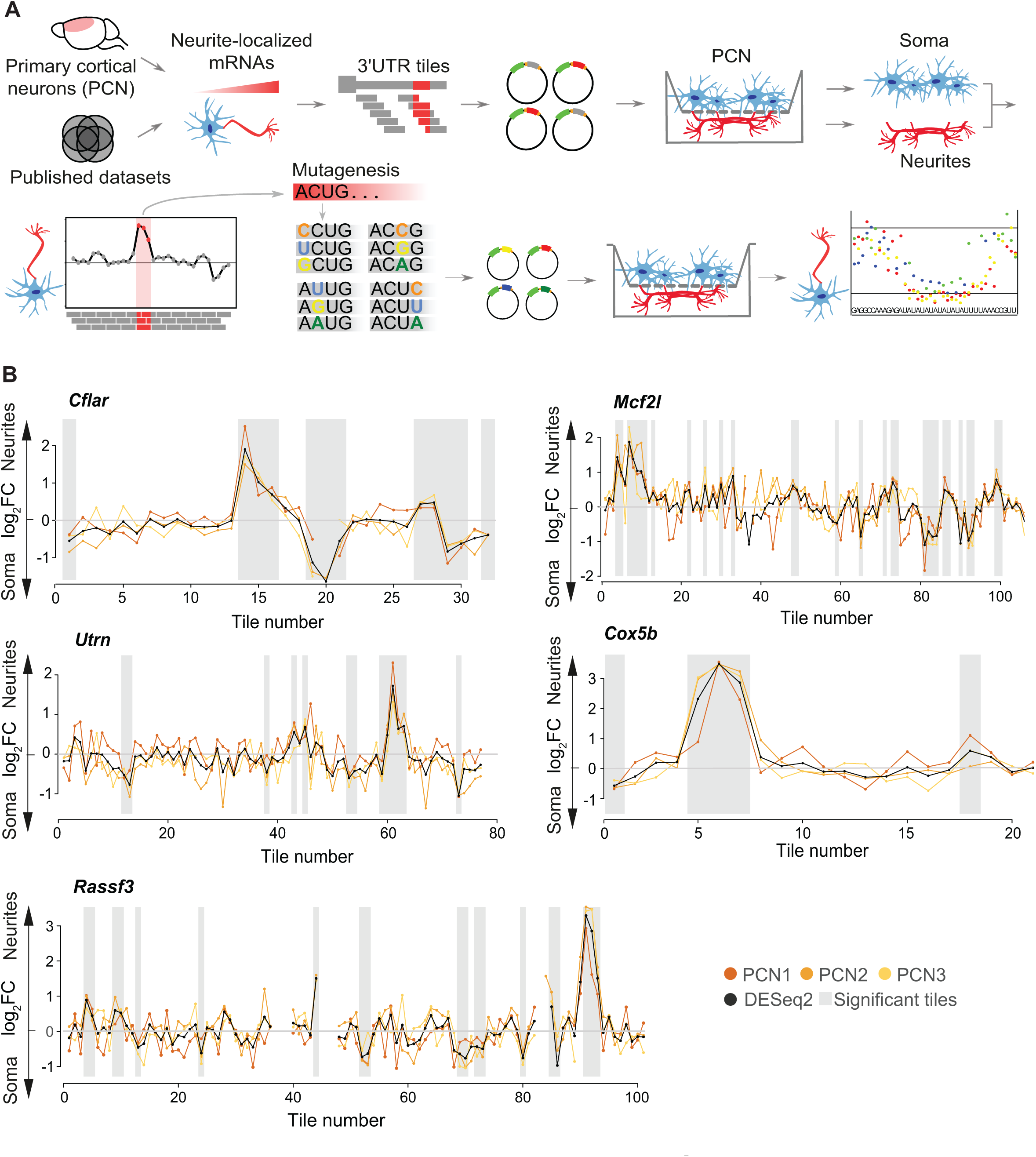
N-zip identifies neuronal zipcodes in primary cortical neurons. (**A**) Scheme of N-zip. Integrative analysis is used to identify a group of transcripts localized to neurites in at least two neuronal localization datasets. Fragments tiled across 3’UTRs of these transcripts (tiles) are generated and cloned in 3’UTRs of lentiviral vector, between adapter sequences, to generated a pooled reporter library. Reporter library is delivered into neurons grown on a microporous filter, separating cell bodies from neurites. Neurite and soma libraries are prepared from 3’UTR fragments, flanked by adaptors, to identify fragments sufficient for neurite localization. Mutagenesis of localized fragments allows to map zipcodes in the secondary N-zip. (**B**) N-zip identifies 3’UTR fragments driving RNA localization to neurites of primary cortical neurons (PCN). Specific examples of identified tiled fragments that mediate localization to neurites. Enrichment of a given tile (log_2_-transformed fold change) between subcellular compartments (Y) is plotted against tiled fragment number (X). Neurite/soma ratios for individual biological replicates (shades of yellow: PCN1, PCN2, PCN3) and computed by DESeq2 based on all replicates (black line) are shown. Shaded regions indicate tiles with significant enrichment (P < 0.05) in one of the subcellular compartments. The gene name is shown above each plot.

As the input for this analysis of zipcodes, we selected 99 transcripts enriched in the neurites (log2-transformed fold change (FC) neurites/soma ≥ 1, adjusted p-value < 0.05, see **Methods**) of mouse primary cortical neurons (**Figure S1A**) and at least one other published dataset generated from primary neurons: dorsal root ganglia, cortical, hippocampal or motor neurons (Briese et al., 2016; Middleton et al., 2019; Minis et al., 2014; Rotem et al., 2017; Taliaferro et al., 2016; Tushev et al., 2018; von Kugelgen and Chekulaeva, 2020). We reasoned that 3’UTRs of these transcripts are likely to contain zipcodes.

To narrow down regions containing potential zipcodes, we designed a pool of 4,813 oligos, 75-110 nt in length, tiled across 3’UTRs of selected neurite-enriched transcripts with 15-25 nt offset (**Table S1**). The oligos, flanked by adapter sequences, were cloned into the 3’UTR of an mRNA coding for GFP, and the resulting pooled lentiviral library was delivered into mouse cortical neurons. To restrict the expression of the library to neurons, we drove the expression of the reporter with a neuron-specific synapsin I promotor. Infected neurons were cultured on a microporous membrane in a way that ensured that soma stayed on top of the membrane and neurites grew through the pores the lower side (Ciolli Mattioli et al., 2019). Subsequently we isolated the neurites and the soma, carried out RT-PCR and prepared amplicon sequencing libraries, referred to as N-zip libraries. The efficiency of the separation between neurites and soma was confirmed by western blotting based on somatic and neuritic markers (**Figure S1B**). Because the reporter library fragments had been flanked with adapters, we could specifically amplify these fragments and exclude endogenous RNAs.

Our analysis of triplicate N-zip libraries identified fragments that exhibited a localization to the neurites of primary cortical neurons (**Figure 1B, Table S1**). For example, we detected a neurite localization (log2FC neurites/soma ≥ 1) of tiled fragments 7-10 derived from the *Mcf2l* mRNA, which encodes the guanine nucleotide exchange factor for CDC42 and RHOA. This protein mediates the formation and stabilization of the glutamatergic synapses of cortical neurons and has been associated with mental retardation and autism (Hayashi et al., 2013). Similarly, we observed a neuritic enrichment of fragment 61 of the *Utrophin* mRNA (*Utrn*), which encodes a component of a dystrophin glycoprotein complex. This complex links the actin cytoskeleton to the extracellular matrix and plays a role in the formation of neuromuscular junctions (reviewed in Belhasan and Akaaboune, 2020).

To fine-map the sequences that mediate localization, we selected fragments from 16 genes that exhibited localization to neurites of primary cortical neurons (selected from genes with mean log2FC neurites/soma ≥ 1 for at least one 3’UTR fragment, adjusted p-value < 0.1, **Table S1**). We performed extensive mutagenesis of these fragments, generating a secondary N-zip library that included 6,266 sequences (**Figure 1A**). In these cases: (1) we introduced every possible single point mutation, (2) we mutated G ↔ C and A ↔ U within 2 nt, 5-nt and 10-nt windows across each fragment (**Table S2**). Our analysis of mutated fragments identified two specific motifs required for their localization to neurites: CUACCUC and (AU)n motifs (**Figure 2**).

**Figure 2.**
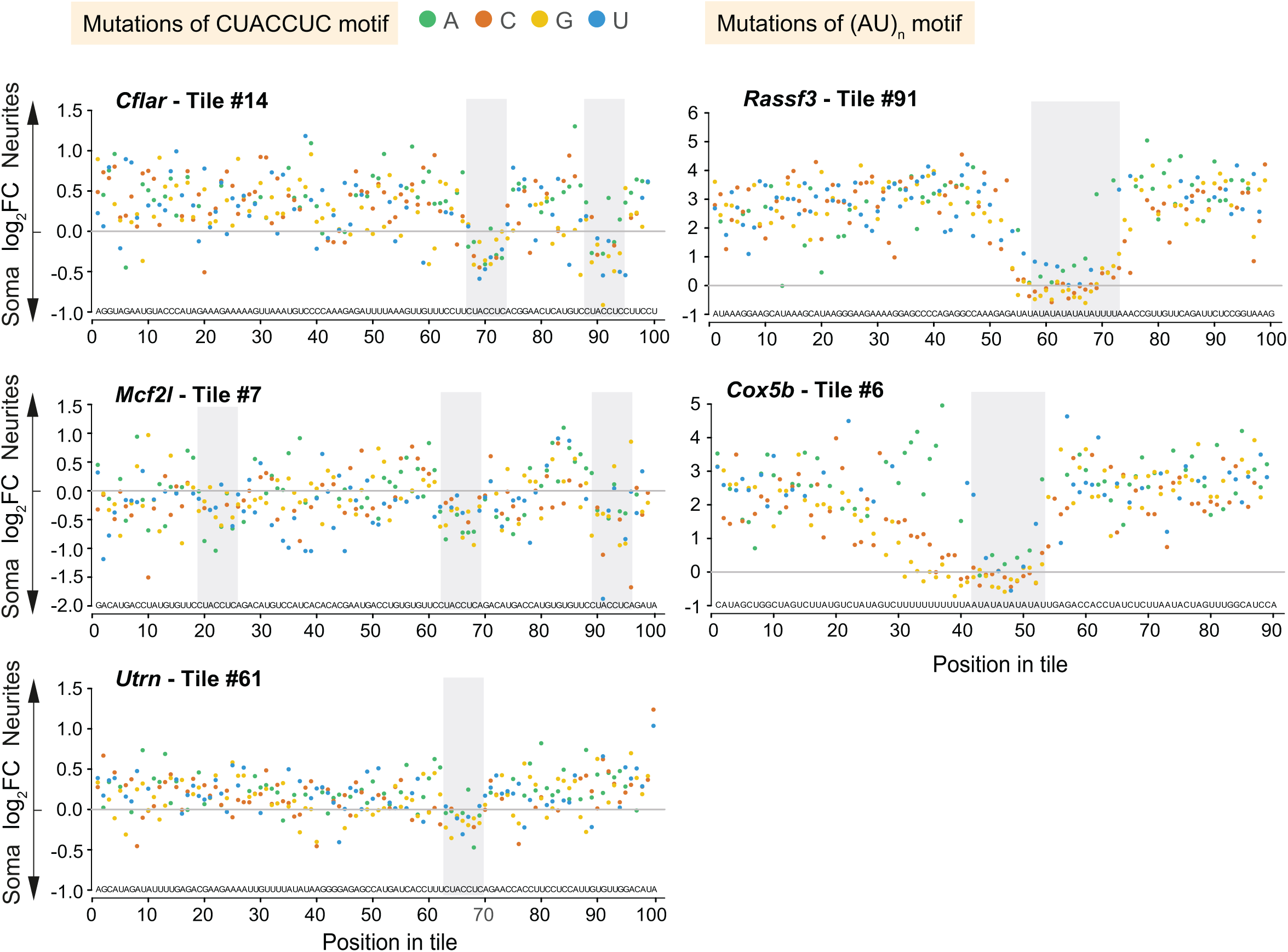
N-zip combined with mutagenesis maps motifs driving mRNA localization to neurites of primary cortical neurons. Selected localized tiled fragments were mutagenized and used for the secondary N-zip in primary cortical neurons as shown in **Figure 1A** (step “Mutagenesis”). Specific examples of mutated motifs that mediate localization to neurites are shown. The data are presented as in **Figure 1B**. The initial sequence of mutagenized fragment is shown above X axis and introduced point mutations are indicated with green (A), orange (C), yellow (G) and blue (U) dots. The gene name and tile number are shown above the plot.

Mutations in CUACCUC shifted the localization of *Cflar*-14, *Mcf2l*-7 and *Utrn*-61 fragments toward the soma. For the (AU)n motif, any mutation of the (AU)8 stretch in *Rassf3-91* dramatically reduced neurite enrichment. Similarly, the *Cox5b-6* (AU)6 motif turned out to be absolutely essential for localization, with contributions from flanking U and A bases and from an additional (U)11 stretch. This latter region could tolerate mutations to A, but not to G or C. Mutations that added (AU) or (UA) repeats to fragments, that had originally exhibited a smaller number of AU or UA repeats, induced neurite localization.

In both the original and the secondary library, (AU)n was associated with neurite localization for n ≥ 5. In the original library, these motifs were also found in *Map2* fragments 29-32, *Ppp1r9b* fragments 56-59, *Shank3* fragments 58-60, and *Tmcc2* fragments 36-39, and some showed conservation in other mammals (**Figure S2A, Table S1**). Curiously, in some cases our mutagenesis introduced CUACCUC and (AU)n stretches into heterologous sequences (*e*.*g. Kif1c, Cald1, Map2*-5, *Camk2n1*-12, **Figure S2B**). These artificially created motifs resulted in the localization of the entire fragment to neurites, showing that these motifs are not only necessary, but also sufficient for localization to neurites.

### Let-7 directs localization of its target mRNAs to neurites

Analysis of miRbase (Kozomara et al., 2019) showed that the CUACCUC motif, which we had identified as a *de novo* zipcode in our N-zip libraries (**Figures 2, S2B**), represents the binding site for the seed of the let-7 miRNA family (**Figure 3A**, top). miRNAs are key regulators of gene expression and target more than half of all genes in mammals (Friedman et al., 2009). They do so by pairing with complementary sites in their target mRNAs; this recruits a complex of proteins which destabilizes the mRNAs (reviewed in Braun et al., 2013; Chekulaeva and Filipowicz, 2009). The miRNA seed is a conserved sequence at positions 2-7 from the miRNA 5’-end which binds to target mRNAs via a perfect complementary base-pairing. Indeed, every point mutation in CUACCUC affected localization of tiles containing this motif (**Figure 2B)**. This is to be expected from a miRNA seed site, but not from a consensus RBP-binding motif. Further, in mouse brain the position of the CUACCUC motif in *Utrn-61* matched the summit of a <u>c</u>ross-linking immuno<u>p</u>recipitation (CLIP) peak for AGO2, which is the core component of the miRNA repression complex (**Figure 3A**, bottom).

**Figure 3.**
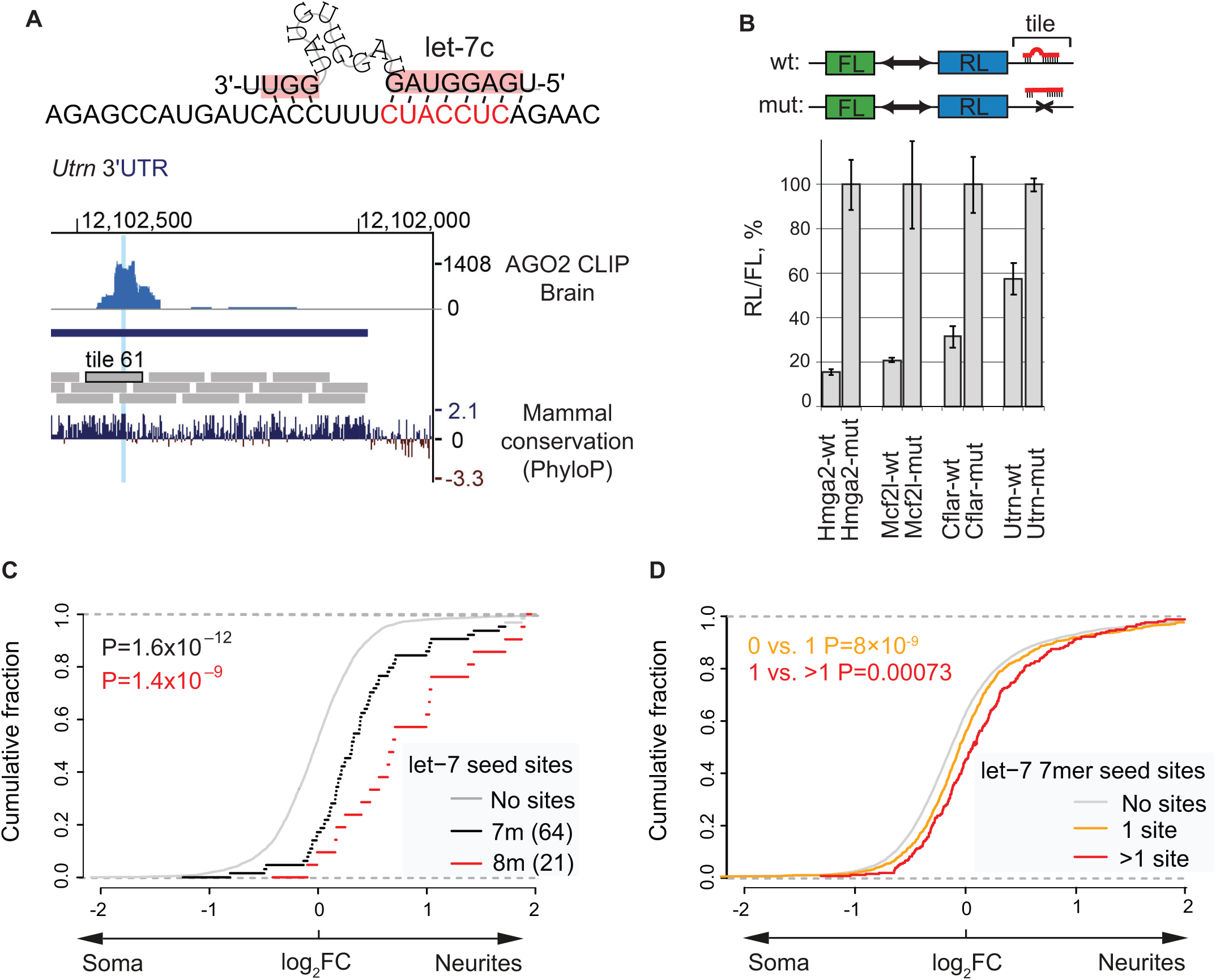
Let-7 binding sites direct mRNA localization to neurites in primary cortical neurons. (**A**) Neurite-localized motif CUACCUC is a binding site for let-7 seed sequence. Top: the scheme showing base-paring between let-7c and Utrn-61 tile as predicted by mfold (Zuker et al., 2003). Bottom: UCSC genome browser view of *Utrn* 3’UTR, showing coverage of AGO2 HITS-CLIP reads in mouse brain (Chi et al., 2009, upper track) around the position of let-7 seed in Utrn-61 identified in N-zip (highlighted in blue). PholyP mammal conversation score (lower track) shows high sequence conservation in vertebrate evolution. (**B**) Validation of functionality of let-7 sites in Cflar-14, Mcf2l-7 and Utrn-61 in luciferase reporter assay. Schematic representation of constructs used in transfection experiments is shown on the top: *Renilla* (RL) and firefly (FL) luciferase are produced from a bi-directional promoter, with indicated N-zip tiles, either wild type (wt) or with mutated let-7 sites (mut), inserted downstream of RL. Indicated luciferase reporters were transfected in HeLa-rtTA cells. As a positive control, analogous reporters with *Hmga2* 3’UTR, either wt (hmga2-wt) or lacking let-7 sites (hmga2-mut), were used. For each reporter pair, RL activity was normalized to that of FL and presented as a percentage of luciferase activity produced in the presence of mutated reporter. Values represent means +/-SD from 3 experiments. (**C**) N-zip reporter mRNAs with let-7 sites are enriched in neurites of primary cortical neurons. Cumulative distribution function (CDF) showing fractions of N-zip mRNAs (Y) plotted against neurite/soma enrichment (X) for transcripts with no let-7 sites (grey), at least one 7mer (black) or 8mer let-7 site (red). The number of detected tiles with indicated let-7 sites is shown in the legend. P-values computed with two-sided Wilcoxon rank-sum test. (**D**) Endogenous mRNAs with let-7 sites are enriched in neurites of primary cortical neurons. The data are plotted as in (**C**), for transcripts with no (grey), one (yellow) or more than one (red) let-7 sites.

To confirm that the let-7 binding sites in *Cflar-14, Mcf2l-7* and *Utrn-61* are functional, we performed a let-7 luciferase reporter assay (**Figure 3B**). As a positive control, we used a reporter bearing the 3’UTR of *Hmga2* gene (Hmga2-wt), a validated let-7 target (Chekulaeva et al., 2011; Hock et al., 2007; Mayr et al., 2007), downstream of the *Renilla* luciferase coding sequence (RL). As a transfection control, we made use of firefly luciferase reporter (FL) without let-7 sites, expressed from the same vector. Endogenously produced in HeLa cells let-7 repressed Hmga2-wt > 5-fold compared to a mutant version lacking let-7 sites (Hmga2-mut, **Figure 3B**). To evaluate the role of let-7 seed sites in *Mcf2l-7, Cflar-14* and *Utrn-61* fragments, we generated analogous luciferase reporters bearing the corresponding tiled fragments in their 3’UTRs (Mcf2l-wt, Cflar-wt and Urtn-wt). As negative controls, we mutated let-7 seeds in the tested regions (Mcf2l-mut, Cflar-mut and Utrn-mut). Compared to the mutated versions, the Mcf2l-wt reporter with three let-7 sites was repressed about 5-fold, the Cflar-wt reporter with two let-7 sites about 4-fold and the *Utrn* reporter with a single let-7 site – about 1.7-fold. These data confirmed that the let-7 sites are functional in the neuritically localized 3’UTR fragments that we tested.

We next wondered whether other let-7 targets are localized to the neurites of primary cortical neurons. To test this, we examined the frequency of let-7 binding sites across differentially localized transcripts in our N-zip libraries. Consistent with a role of let-7 binding sites in mRNA localization, we observed an enrichment of transcripts bearing let-7 7mer seeds in neurites (black line, **Figure 3C, Table S1**) compared to transcripts without let-7 seeds (grey line). The effect was even stronger for extended 8mer seeds (red line).

We next wondered whether let-7 sites also contribute to the localization of endogenous mRNAs. To test this, we analyzed the frequency of let-7 binding sites in differentially localized endogenous mRNAs. We detected an enrichment of let-7 site-bearing transcripts in neurites, the degree of which depended on the number of let-7 sites (grey line: no sites, yellow line: one site, red line > 1 site, **Figure 3D, Table S3**). This let-7 site-dependent shift in localization was less profound for endogenous mRNAs than for N-zip mRNA reporters, probably because other regions in endogenous full-length 3’UTRs contribute to their localization in a combinatorial manner.

Our global analysis of miRNA seeds in N-zip libraries revealed that let-7 binding sites were enriched compared to sites of other miRNAs in neurites (**Figure 4A**). To investigate the potential underpinnings of this specificity, we used small RNA-seq to analyze miRNA expression levels in cortical neurons. We found that let-7 is the most abundant miRNA in cortical neurons (> 30 % of all miRNA reads, **Figure 4B, Table S4**), providing an explanation for the preferential influence of let-7 on the neurite-enriched transcriptome. Given the role of miRNAs in mRNA stability, we compared levels of N-zip reporters that contained let-7 seeds with those in which let-7 seeds had been mutated, both in soma and neurites. We observed that mutations of let-7 binding sites stabilized N-zip reporter mRNAs (**Figure 4C**, compare red and grey boxes), and this effect was stronger in soma than in neurites. These data suggest that let-7 promotes the enrichment of its targets in neurites by destabilizing them more potently in the soma.

**Figure 4.**
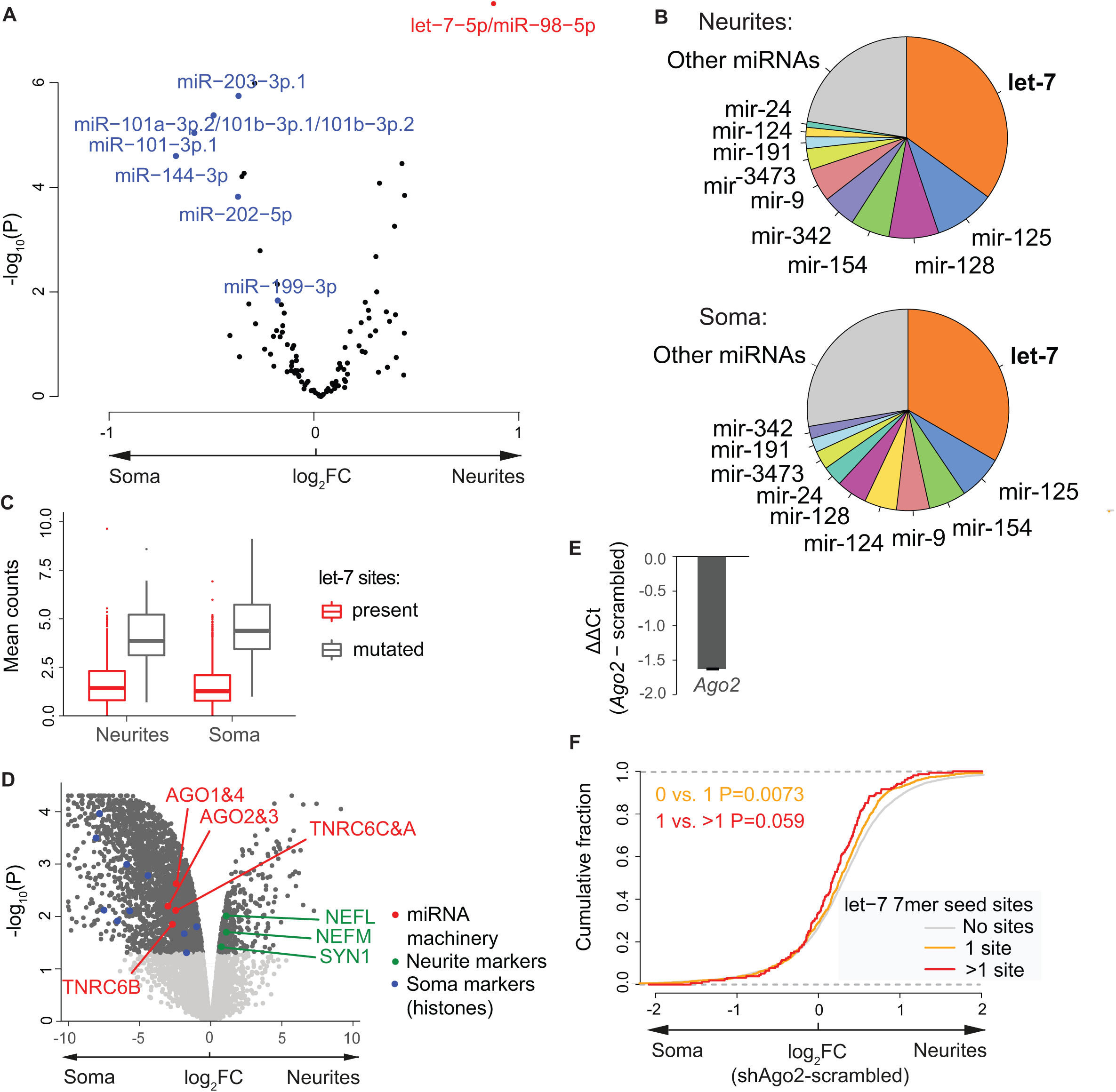
Let-7 functions by preferentially destabilizing its target mRNAs in soma of primary cortical neurons. (**A**) Let-7 binding sites are enriched in neurite-localized mRNAs in primary cortical neurons. Volcano plot showing the mean neurite/soma enrichment (log2 fold change) of tiles containing 7mer matches to the seeds of individual miRNA families in the N-zip mRNA reporters. miRNAs with statistically significant enrichment in one of the compartments (P < 0.05, computed by two-sided t-test) are labeled in red (neurite-enriched) and blue (soma-enriched). (**B**) Let-7 is the most abundant miRNA in primary cortical neurons. Pie-charts showing the percentage of different miRNAs in neurites and soma of primary cortical neurons, determined by small RNA-seq. (**C**) Mutations of let-7 binding sites lead to stronger increase of N-zip mRNA reporter levels in soma. Boxplots of mean (n=3) normalized counts from either soma or neurite mutated N-zip libraries. Reporter tiles were separted into groups based on occurrence of let-7 seed binding site (CUACCUC, red: let-7 sites are preserved, grey: let-7 site is mutated). (**D**) Proteomic analysis of isolated soma and neurites indicates somatic enrichment of the core components of the miRNA repression complex (AGO1, 2,3 and 4; TNRC6A, B and C, red). Volcano plot showing -log10 P-values (Y) plotted against log2 fold change of iBAQ values, normalized to GAPDH in each compartment. Significant values (FDR <5%) are shown in dark grey, non-significant ones in light grey. Neurofilaments (NEFL, NEFM) and Synapsin (SYN1) are used as neurite markers (green), and histones – as soma markers (blue). (**E**) RT-qPCR showing the efficiency of *Ago2* depletion with shRNA in primary cortical neurons. Difference in Ago2 expression levels (ΔΔCt) between *Ago2*-depleted and control scrambled shRNA samples is plotted in Y axis. *Gapdh* was used as normalization control. Values represent means +/-SD from 3 biological replicates. (**F**) Ago2 knockdown in primary cortical neurons shifts let-7 targets towards soma. CDF showing fractions of endogenous mRNAs with no let-7 sites (grey), one (yellow) and more let-7 site (red), as measured by mRNA-seq (Y), plotted against changes in neurite/soma enrichment upon *Ago2* knockdown (X). P-value computed using two-sided Wilcoxon rank-sum test.

Because levels of let-7 were comparable in neurites and the soma, we next used proteomics to examine the expression levels of miRNA repression complex components. This analysis showed that the core components of the miRNA repression complex, including AGO and TNRC6 family members, are enriched in soma (**Figure 4D**, red, **Table S5**). We controlled the efficiency of the separation of soma and neurites by measuring neuritic (neurofilaments and synapsin, green) and somatic (histones, blue) markers. In summary, these data explain a higher activity of let-7 activity in the soma, which ultimately leads to an enrichment of let-7 targets in neurites.

To experimentally confirm the role of the miRNA pathway and let-7 in mRNA localization, we depleted *Ago2*, the key component of the miRNA repression complex, in primary cortical neurons using shRNA. RT-qPCR analysis confirmed a ∼ 70% depletion of *Ago2* compared to the controls transduced with a scrambled shRNA (**Figure 4E, Table S3**). We followed this up with an isolation of neurites and soma and carried out an mRNA-seq analysis of *Ago2*-deleted and control samples. We next analyzed how the enrichment of let-7 targets in neurites changes upon *Ago2* depletion. When compared to mRNAs that did not contain let-7 sites (grey line, **Figure 4F**), let-7 targets shifted towards the soma, an effect whose degree depended on the number of sites in the transcript (yellow line: 1 let-7 7mer seed match, red line: > 1 let-7 7mer seed match).

*Ago2* depletion affects the entire miRNA pathway. To examine the specific effects of let-7 on mRNA localization, we depleted let-7 with a sponge construct bearing six binding sites for let-7 in its 3’UTR (Kumar et al., 2008, **Figure S3A**). To show that the sponge construct can indeed alleviate silencing by let-7, we performed a let-7 reporter assay in HeLa-rtTA cells as described in **Figure 3B**. Let-7 efficiently repressed Hmga2-wt mRNA as compared to the mutant reporter from which let-7-binding sites had been deleted (compare Hmga2-wt with Hmga2-mut, green outlined bars, **Figure S3A**). When we depleted endogenous let-7 with increasing amounts of our GFP-let-7 sponge-encoding construct (red outlined bars), this alleviated the repression of the Hmga2-wt reporter. A negative control, consisting of a GFP-encoding construct without let-7 sponge sites, did not interfere with repression (green outlined bars).

Having established the functionality of the GFP-let-7 sponge, we used it to deplete let-7 in primary cortical neurons grown on a microporous membrane. The GFP-encoding construct was used as a negative control in place of the sponge. We then analyzed changes in mRNA localization by comparing GFP-let-7 and control GFP samples. We observed that the depletion of let-7 shifted the localization of let-7 targets towards the soma (**Figure S3B, Table S3**). These data validated the effect of let-7 on localization of endogenous mRNAs in primary cortical neurons.

### (AU)_n_ motif-containing mRNAs recruit HSB1L protein to localize to neurites

Our N-zip analysis identified the (AU)_n_ motif as a *de novo* zipcode that mediates mRNA localization to the neurites of primary cortical neurons. We next analyzed how the length of the (AU)_n_ stretch affected mRNA localization in N-zip libraries. We found that mRNAs with six or more AU repeats were enriched in neurites (**Figure 5A, Table S1**). We then examined how the (AU)_n_ motif affected the localization of endogenous mRNAs. We observed that transcripts with six or more AU repeats in their 3’UTRs (red line, **Figure 5B, Table S3**) were shifted to neurites compared to transcripts with fewer than six AU repeats or none (grey line). The effect was weaker for endogenous transcripts than for N-zip mRNA reporters with short 3’UTRs, presumably due to other regulatory sequences in the full-length 3’UTRs. To test whether the (AU)_n_ motif plays a role in mRNA localization in other cell types, we analyzed the effects of the length of the (AU)_n_ stretch in the neuroblastoma cell lines CAD and N2a (Mikl et al., 2021). Strikingly, transcripts with (AU)_n_ with n ≥ 10 were consistently enriched in the neurites of both cell lines (**Figure S4A**), suggesting that the role of this motif in mRNA localization is conserved.

**Figure 5.**
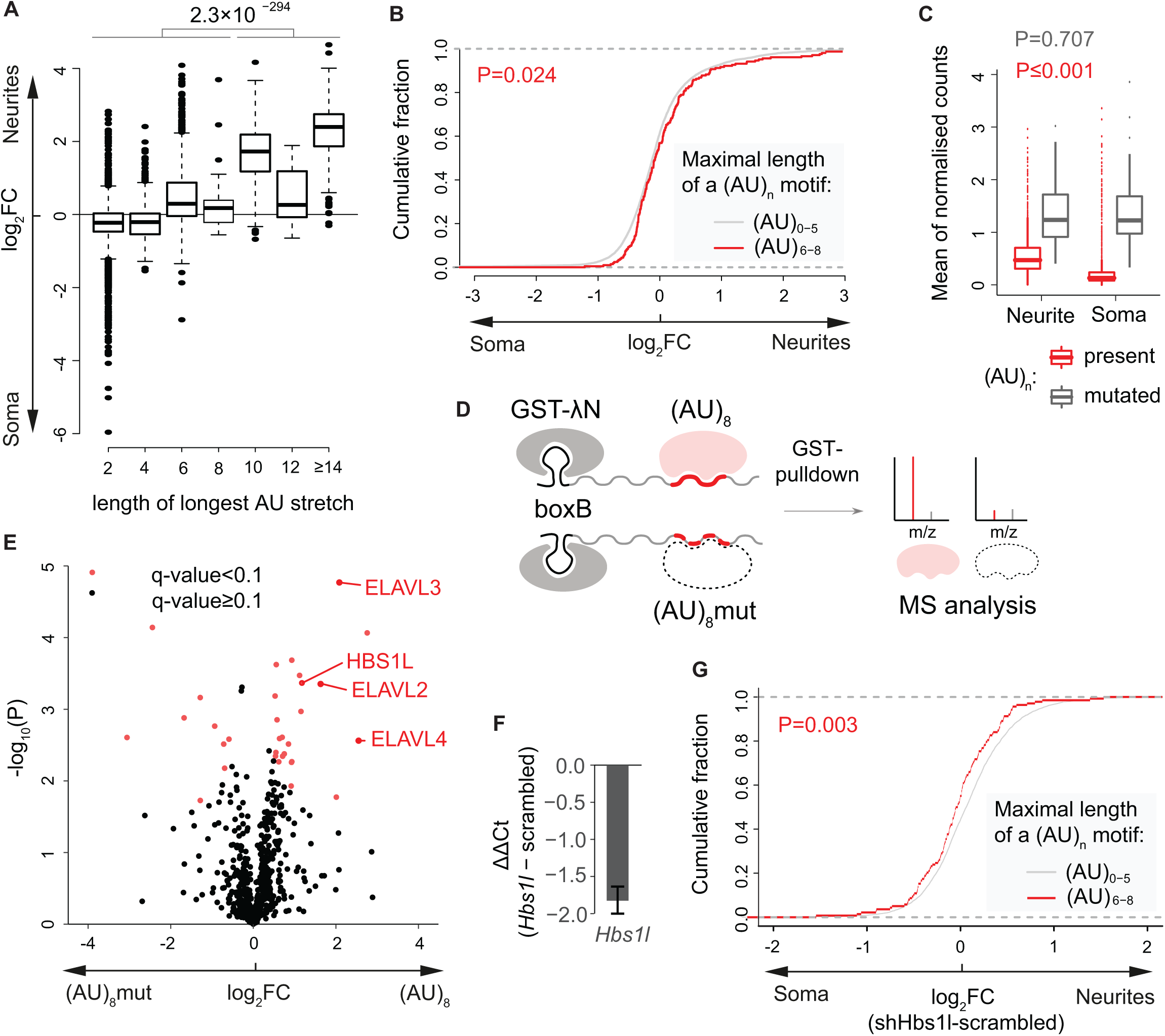
HBS1L binds (AU)_n_ motif to directs mRNA localization to neurites of primary cortical neurons. (**A**) Boxplot showing neurite/soma enrichment as a function of the maximal length of the (AU)_n_ stretch length in the tiles in N-zip mRNA libraries. (**B**) CDF showing fractions of endogenous mRNAs with (AU)_0-5_ (grey) and (AU)_6-8_ stretches (red), as measured by mRNA-seq (Y), plotted against neurite/soma enrichment. P-value computed using two-sided Wilcoxon rank-sum test. (**C**) Mutations of (AU)_n_ leads to stronger increase of N-zip mRNA reporter levels in soma. Boxplots of mean (n=3) normalized counts from either soma or neurite mutated N-zip libraries. Reporter tiles were separted into groups based on occurrence of (AU)_n_ (red: (AU)_n_ is preserved, grey: (AU)_n_ is mutated). (**D**) Scheme for GRNA chromatography in combination with mass spectrometry to identify RBPs bound to (AU)_8_-containing RNA. BoxB-tagged (AU)_8_-carrying mRNA was immobilized on N-GST-glutathione beads and incubated with mouse brain protein lysates. Bound RBPs were eluted with RNAse A and analyzed by mass spectrometry. RNA with mutated (AU)_8_ was used as a negative control. (**E**) (AU)_n_ motif binds HBS1L and nELAVL proteins. Volcano plot showing proteins enriched in (AU)_8_-GRNA chromatography. -log_10_ P-values (Y, Student’s two sample t-test with equal variance) are plotted against log_2_ fold change of LFQ (label free quantification) values between (AU)_8_ and mutant RNA pulldowns (X). Proteins with significant q-value (FDR <5% using BH correction) are marked red. (**F**) RT-qPCR showing the efficiency of *Hbs1l* depletion with shRNA in primary cortical neurons. Difference in *Hbs1l* expression levels (ΔΔCt) between Hbs1l-depleted and control scrambled shRNA samples is plotted in Y axis. Gapdh was used as normalization control. Values represent means +/-SD from 3 biological replicates. (**G**) Depletion of *Hbs1l* in primary cortical neurons shifts (AU)_n_-containing mRNAs towards soma. CDF showing fractions of endogenous mRNAs with no or short (AU)_n_ stretch (n=0-5, grey) and a long (AU)_n_ stretch (n > 5, red), as measured by mRNA-seq (Y), plotted against changes in neurite/soma enrichment upon *Hbs1l* depletion (X). P-value computed using two-sided Wilcoxon rank-sum test.

We next examined how (AU)_n_ affects transcript levels in soma and neurites. Mutations of (AU)_n_ stabilized N-zip reporter mRNAs (**Figure 5C**, compare red and grey boxes), and this effect was stronger in the soma than in neurites. These data suggest a mechanism similar to that we described above for the let-7 binding sites: (AU)_n_ mediates enrichment of transcripts in neurites through their preferential destabilization in the soma.

The localization of zipcodes is mediated through binding to RBPs which recruit components of the transport machinery or regulators of mRNA stability (reviewed in Glock et al., 2017). We decided to investigate the RBPs that are recruited by the newly identified (AU)_n_ motif and might mediate its localization. To achieve this, we used GRNA chromatography (Czaplinski et al., 2005), an RNA affinity capture approach, combined with mass spectrometry (**Figure 5D**). We tagged the (AU)_8_-containing *Rassf3*-91 fragment with five copies of the boxB sequence, which has strong affinity to lambda N peptide, and incubated the resulting (AU)_8_-boxB and negative control with mutated motif (AU)_8_mut-boxB with cortical lysates. We then isolated the complexes formed on (AU)_8_-boxB and (AU)_8_mut-boxB RNAs using lambda N-GST fusion protein immobilized on glutathione beads. Protein binders, eluted with RNAse A, were used for further proteomic analysis.

Using this approach, we identified 23 proteins as significantly enriched in (AU)_8_ versus (AU)_8_mut complexes (FDR 5%, **Figure 5E, Table S6**). Among them, were <u>HBS1</u>-like protein (HBS1L), critical for cerebellar neurogenesis (Terrey et al., 2021), and <u>n</u>euronal members of the <u>ELAV</u>-like (nELAVL) family (ELAVL2/HuB, ELAVL3/HuC and ELAVL4/HuD) with the roles in neuronal differentiation, learning and memory (DeBoer et al., 2014; Quattrone et al., 2001; Vanevski and Xu, 2015).

HBS1L and nELAVLs were reported to have opposite effects on mRNA fate: HBS1L is an mRNA decay factor (Kalisiak et al., 2017; Simms et al., 2017), while nELAVL proteins stabilize bound mRNAs by preventing their association with destabilizing proteins (reviewed in Mirisis and Carew, 2019). As localization of (AU)_n_-containing mRNAs to neurites is due to their preferential destabilization in soma (**Figure 5C**), HBS1L is a plausible player in this mechanism. Thus, we depleted *Hbs1l* with an shRNA (**Figure 5F**), and examined how this depletion affected localization of transcripts with (AU)_n_ motif by mRNA-seq of isolated neurites and soma. Remarkably, *Hbs1l* depletion shifted the localization of transcripts with a long (AU)_n_ motif ((AU)_6-8_, red line, **Figure 5G**, see also **Figure S4B**) towards the soma, when compared with transcripts with a short or no (AU)_n_ motif ((AU)_0-5_, grey line). In contrast, the same approach applied to nELAVL-depleted neurons (**Figure S4C**) showed no correlation between the length of (AU)_n_ and changes in mRNA localization (**Figure S4D**). Similarly, depletion of other proteins detected in (AU)_8_-GRNA chromatography - RC3H2, MAST1 and ZFP871 - did not interfere with the localization of (AU)_n_-containing mRNAs (**Table S6** and data not shown). Thus, these data confirmed the role of HBS1L in localization of (AU)_n_-containing transcripts in primary cortical neurons.

## DISCUSSION

mRNA localization has many advantages over the localization of proteins as a mechanism of generating high concentrations of specific proteins at defined sites in cells: (1) ectopic protein activity is avoided; (2) the local proteome can change quickly in response to specific local stimuli (*e*.*g*. neurotransmitters and growth factors); (3) economy: each localized mRNA can be used to produce many localized proteins; (4) mRNAs can be targeted to distinct subcellular localizations based on zipcode information in their 3’UTRs, without changing the structure or functions of the proteins they encode. Our prior work has shown that at least half of the neurite-localized proteome can be accounted for by the localization of mRNAs to these regions, which highlights the crucial importance of asymmetrical mRNA localization in neurons (Zappulo et al., 2017 6881).

As well as transporting mRNAs along cytoskeletal fibers with the help of motor proteins, localization can be achieved by degrading mRNAs in other regions where they should not be on hand. This latter mechanism has been described for mRNAs of *Hsp83* and *nanos*, for example. Their mRNAs are degraded throughout *Drosophila* eggs via the Smaug-mediated recruitment of the CCR4-NOT deadenylation complex, but remain stable at the posterior pole (Chen et al., 2014; Ding et al., 1993; Semotok et al., 2005). Prior studies have reported a local processing of miRNAs (Sambandan et al., 2017) and a modification of the components of the miRNA pathway in response to synaptic activity (Ashraf et al., 2006; Banerjee et al., 2009; Muddashetty et al., 2011). However, to the best of our knowledge, a localization-dependent degradation of mRNAs by miRNAs has not been previously described. Given the known role of miRNAs in mRNA degradation (reviewed in Braun et al., 2013; Chekulaeva and Filipowicz, 2009; Fabian and Sonenberg, 2012), it seems highly likely that this could be used as a mechanism to establish pools of specific mRNAs in some areas of the cell by degrading them in others. Our N-zip analysis identified the binding site for a miRNA – let-7 – as a zipcode (**Figures 1B and 2**). Let-7 is the most highly expressed miRNA in the mammalian brain (Lagos-Quintana et al., 2002; Petri et al., 2017; Sambandan et al., 2017) (**Figure 4B**), and is involved in neuronal differentiation (Schwamborn et al., 2009), regeneration (Li et al., 2015; Zou et al., 2013), and synapse formation (Caygill and Johnston, 2008; Edbauer et al., 2010). Our results point to a new neuronal function for let-7 in mediating the localization of mRNAs.

Intriguingly, let-7 was equally abundant in the neurites and soma of primary cortical neurons (**Figure 4B**), raising the question about the mechanism underpinning the higher let-7 activity in soma. One explanation came from our proteomic analysis, which has shown that protein components of miRNA machinery are enriched in soma (**Figure 4D**). This suggests that a constellation of factors contribute to the localization of specific mRNAs in neurites: the number of miRNA binding sites in a particular 3’UTR, the presence of miRNAs, and quantities of components that bind and degrade them. Any bias in these elements can lead to a preferential localization of mRNAs.

We identified the (AU)_n_ motif (n > 5) as another *de novo* zipcode in primary cortical neurons. (AU)_n_ is found in a number of important neuritically-enriched mRNAs, including *Map2* and *Shank3* (**Table S1, Figure S2A**). RNA affinity capture showed that the (AU)_n_ motif is bound by an mRNA decay factor HBS1L and nELAVL family proteins (**Figure 5D**), both having important functions in neuronal development (DeBoer et al., 2014; Quattrone et al., 2001; Terrey et al., 2021; Vanevski and Xu, 2015). Based on RIP-chip and SELEX experiments (Cook et al., 2011), ELAVL proteins bind AU-rich elements (ARE), including UUUAUUU and some of its variations. AREs bear some resemblance to (AU)_n_ motif, which may explain detection of nELAVLs among (AU)_n_ interactors in our GRNA chromatography. However, ELAVL proteins stabilize their bound mRNAs (reviewed in Mirisis and Carew, 2019), while our analysis showed that localization of (AU)_n_-containing transcripts is mediated by selective mRNA destabilization (**Figure 5C**). Indeed, depletion of nELAVLs did not decrease neurite enrichment of (AU)_n_-containing transcripts (**Figure S4D**).

Our depletion experiments showed that HBS1L mediated localization of mRNAs with the (AU)_n_ motif (**Figure 5G**). HBS1L is involved in mRNA quality control pathways, including No-Go decay (NGD) and Nonstop decay (NSD), that degrade mRNAs with stalls in translation elongation (reviewed in Doma and Parker, 2007). HBS1L belong to GTPases family and is homologous to translation elongation and termination factors eEF1 and eRF3. Here, we uncover a novel role of HBS1L in localization of (AU)_n_-containing transcripts via selective degradation in soma. It remains to be understood how it is recruited to (AU)_n_ motifs to trigger mRNA degradation. Importantly, HBS1L has crucial functions in neuronal development and its deletion lead to cerebellar abnormalities (Terrey et al., 2021).

Curiously, we found that the (AU)_n_ motif is linked to mRNA localization not only in cortical neurons, but also in neuroblastoma cell lines (**Figure S4A**). In contrast, let-7 binding sites function as zipcodes in primary cortical neurons (**Figure 3C-D**), but not in neuroblastoma lines (data not shown). These discrepancies are likely due to differences in the expression of *trans-*acting factors recruited by zipcodes, such as let-7, between primary neurons and neuronal cell lines (Cherone et al., 2019). This example illustrates the merits of using primary cells to identify functional elements with biological roles *in vivo*.

Massively parallel reporter assays (MRPA) have been used to identify elements that regulate mRNA stability, translation and nuclear export (Lubelsky and Ulitsky, 2018; Mishima and Tomari, 2016; Rabani et al., 2017; Sample et al., 2019; Yartseva et al., 2017). Here we expand the potential of this approach with the development of the MRPA-based method N-zip (**Figure 1A**) to identify zipcodes involved in subcellular mRNA localization in neurons. Our analysis identifies two new short motifs, let-7 binding site and (AU)_n_, with a role in mRNA localization in primary cortical neurons.

These motifs help localize both exogenously introduced N-zip mRNA reporters and endogenous mRNAs, but the effect is stronger for N-zip mRNAs reporters. The likely reason is that N-zip reporters have shorter 3’UTR (≤ 150 nt). The full-length 3’UTRs of endogenous mRNAs are longer and thus potentially carry additional regulatory elements that may exercise a finer control over mRNA stability, translation and localization. The ultimate localization of these mRNAs is a combinatorial result based on these multiple regulatory sequences. In support of this, our examination of tiles derived from the same gene shows that a single 3’UTR often harbors sequences that promote and inhibit neurite enrichment (**Figure 1B**). This means that our approach, based on MRPA with short fragments tiled across the 3’UTR of localized mRNAs, can be used to identify zipcodes that would not otherwise be readily detectable in full-length mRNAs.

Decoding the combinatorial effects that actually regulate mRNA localization will require zooming in on shorter sequences with specific functions, such as serving as targets for miRNAs or binding to components of the RNA degradation or transport machinery. Our combination of N-zip with the extensive mutagenesis of localized fragments (**Figure 1A**) permits such a high-resolution mapping of specific motifs that are required for localization.

Such motifs are thought normally to function by recruiting *trans-*acting factors, such as RBPs that regulate their transport or stability. Ideally, identifying such factors can be achieved through RNA affinity capture, either GRNA chromatography used in the current study (**Figure 5E**) or using biotinylated oligos, in combination with mass spectrometry. This combination of techniques can be extended to diverse types of neurons to observe molecular processes in contexts such as healthy development, learning and neurological diseases. It can also be used to study other polarized cell types such as fibroblasts, cancer cells and oocytes. In summary, our work establishes a method for an unbiased, transcriptome-wide identification of zipcodes and zipcode-binding proteins. This is a crucial step toward unraveling how a finite number of patterns produce many types of polarized cells and help them adapt to challenges and changes.

## STAR METHODS

### Key resources table

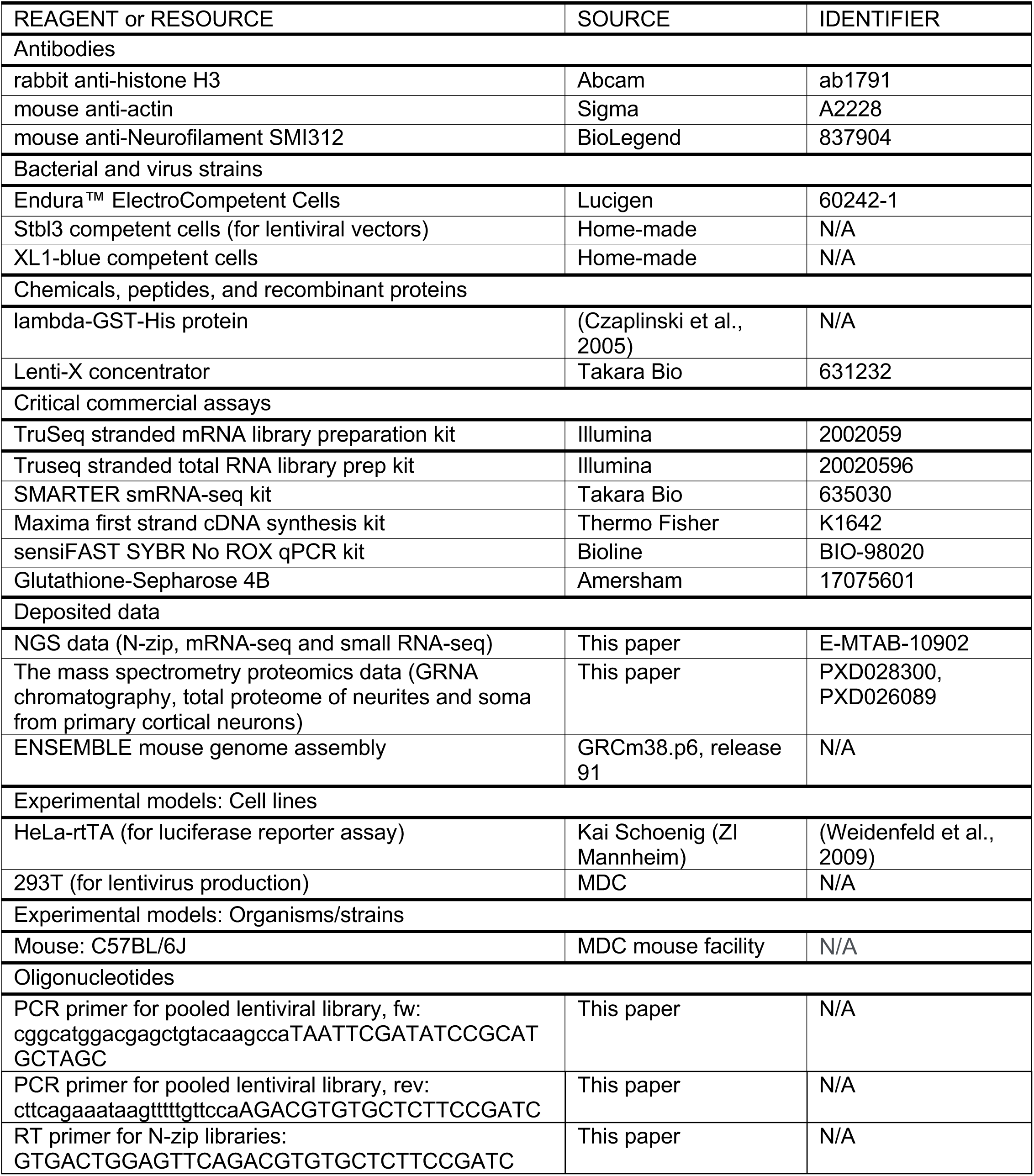
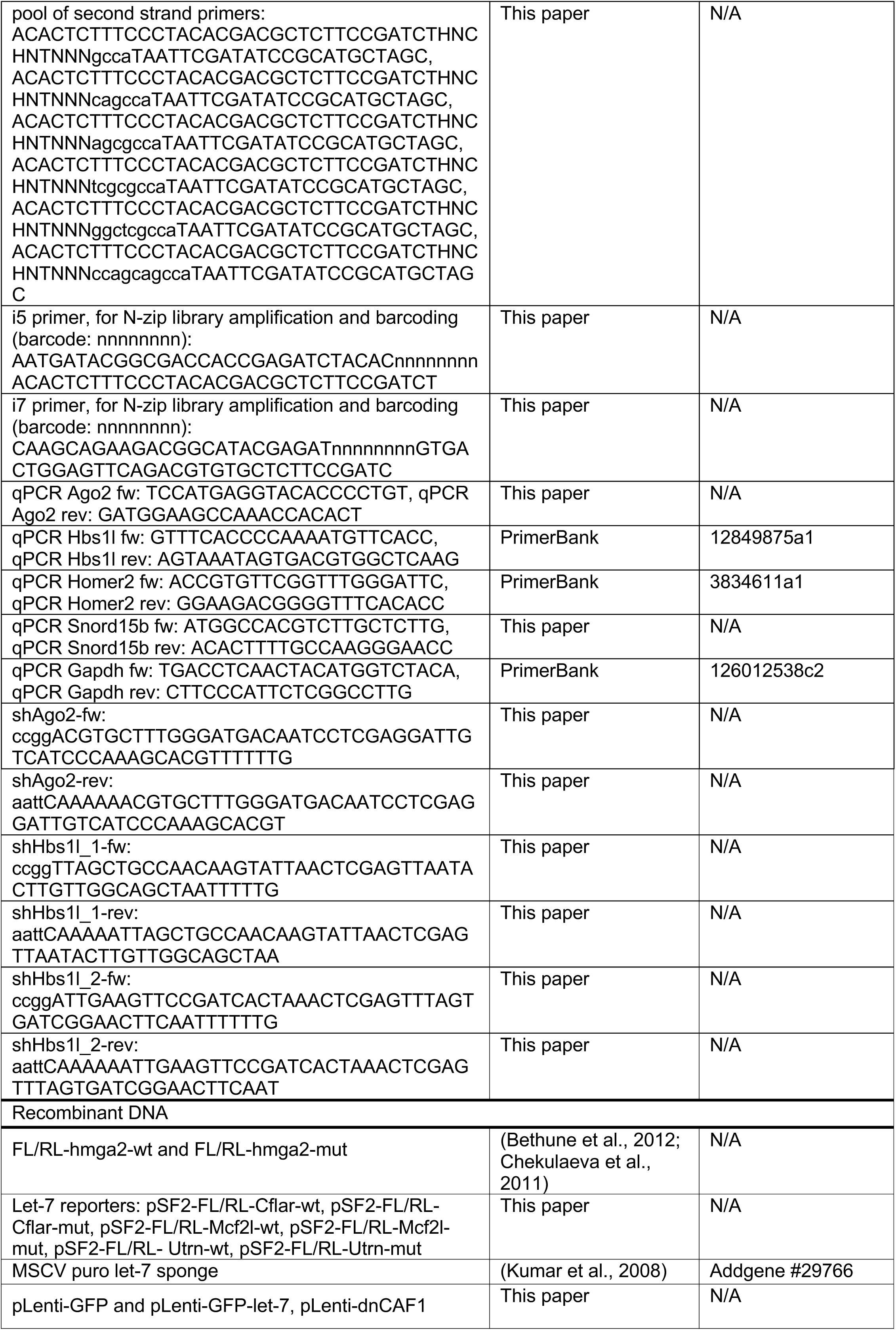
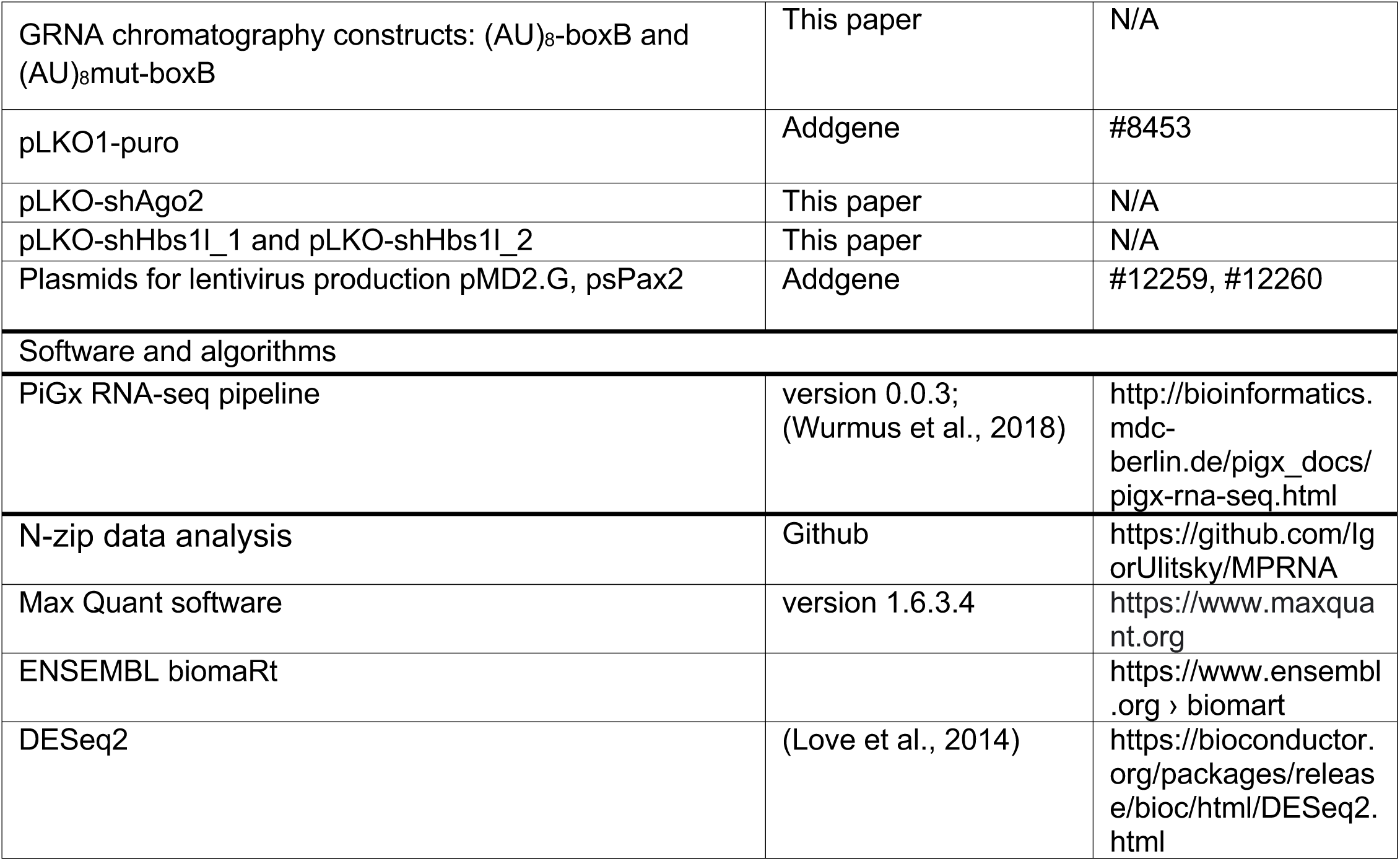

## RESOURCE AVAILABILITY

### Lead contact

Further information and requests for resources and reagents should be directed to and will be fulfilled by the lead contact, Marina Chekulaeva.

### Materials availability

All unique reagents generated in this study are available from the Lead Contact without restriction, or with a Materials Transfer Agreement.

### Data and code availability

NGS data have deposited at the ArrayExpress (accession E-MTAB-10902). The mass spectrometry proteomics data have been deposited to the ProteomeXchange Consortium via the PRIDE (Perez-Riverol et al., 2019) partner repository with the dataset identifiers PXD028300 and PXD026089. The code for N-zip data analysis is available at https://github.com/IgorUlitsky/MPRNA.

## EXPERIMENTAL MODEL AND SUBJECT DETAILS

Primary cortical neurons were isolated from E14 and P0 pups and cultured as previously described (Kaech and Banker, 2006).

HeLa-rtTA cells expressing reverse tetracycline-controlled transactivator (Weidenfeld et al., 2009) and 293T cells, used for lentiviral production, were grown in Dulbecco’s modified Eagle’s medium with GlutaMAX™ supplement (DMEM+ GlutaMAX, GIBCO) with 10% FBS.

## METHOD DETAILS

### Primary cortical neuron culture and lentiviral transduction

Primary cortical neuron culture, separation on soma and neurites and preparation of lentivirus were done as described previously (Ciolli Mattioli et al., 2019). In short, cortical neurons were isolated from E14 and P0 pups and cultured as previously described (Kaech and Banker, 2006). Lentiviral particles were concentrated using Lenti-X concentrator (631232 Takara Bio) to a final volume 200 μl. Virus was applied on cortical neurons between DIV5 and cells were collected at DIV9. For depletion experiments, 25 μl of concentrated virus was added per 10^6^ primary cortical neurons growing on a Millicell cell insert (6-well); for N-zip libraries, 30 μl of concentrated virus was used in combination with depletion construct.

### Luciferase reporter assays

Human HeLa-rtTA cells expressing reverse tetracycline-controlled transactivator (Weidenfeld et al., 2009) were used in luciferase reporter assay. Transfections were done in 96-well plates with polyethylenimine (PEI) using a 1:3 ratio of DNA:PEI. Cells were transfected with 1-3 ng of FL/RL doxycycline-inducible let7 reporter per well. Increasing amounts of GFP-let-7 sponge (40, 60, 80, 100, 125 ng per well) were co-transfected, where indicated. GFP-encoding plasmid was used as a filler, to top up each transfection to the same total amount of DNA. Expression of luciferase reporters was induced with doxycycline (1 μg/ml) and cells were lyzed 24 hr post-transfection. Luciferase activities were measured with a homemade luciferase reporter assay system as described earlier (Mauri et al., 2016).

### DNA constructs

FL/RL-hmga2-wt and FL/RL-hmga2-mut have been previously described (Bethune et al., 2012; Chekulaeva et al., 2011). For other let-7 reporters, we modified the same backbone with bi-directional promoter for simultaneous expression of two genes pSF2.GFPLuc (Loew et al., 2006). First, *Renilla* luciferase was PCR-amplified and cloned into EcoRI/NotI-cut pSF2.GFPLuc, to substitute GFP with RL and produce pSF2-FL/RL. Next, a fragment containing the polyadenylation signal was cloned into NotI site downstream of RL, to generate pSF2-FL/RL-pA. Finally, synthetic oligos corresponding to Cflar-14, Mcf2l-7 and Utrn-61 tiles (**Table S1**) or their mutated versions (CTACCTC → CCATCCC, CTACCTC → GATGGAG) were annealed and cloned between AgeI and NotI sites of pSF2-FL/RL-pA.

To generate GFP-encoding plasmid, GFP ORF was PCR-amplified and cloned between AgeI and EcoRI sited of lentiviral vector with synapsin I promoter (Addgene #20945). During this cloning AgeI site was destroyed and a new AgeI was introduced on a PCR primer downstream of GFP. Resulting pLenti-GFP construct was used to produce GFP-let-7 sponge and pooled N-zip lentiviral libraries. To generate GFP-let-7 sponge, a region bearing six let-7 binding sites was PCR-amplified from MSCV puro let-7 sponge (Addgene #29766) and cloned between AgeI and EcoRI sites downstream of GFP.

To produce (AU)_8_-boxB and (AU)_8_mut-boxB constructs for GRNA chromatography, synthetic oligos corresponding to *Rassf3-91* tile or its mutated version (ATATATATATATATAT → GTACATACATGTACAT) were annealed and cloned between KpnI and NheI sites of pBS-Luc-boxB (Chekulaeva et al., 2006).

pLKO-shAgo2 and pLKO-shHbs1l were generated by cloning of annealed oligos (**Key resources table**) into AgeI and EcoRI-cut pLKO1-puro vector (Addgene #8453).

### Western blotting

Preparation of protein lysates from neurites and soma of primary cortical neurons was done as previously described (Ciolli Mattioli et al., 2019). 5 μg of total protein was separated on a 10 % Laemmli PAAG. Proteins were transferred to the PVDF membrane and analyzed by western blotting. The following primary antibodies were used: rabbit rabbit anti-histone H3 1:5000 (ab1791 Abcam), mouse anti-actin 1:4000 (Sigma A2228), mouse anti-Neurofilament SMI312 1:10000 (837904 BioLegend).

### RNA Extraction and qRT-PCR

For RT-qPCR analysis, RNA was isolated with Trizol (Thermo Fisher), treated with RQ1 DNase I, and reverse-transcribed using the Maxima first strand cDNA synthesis kit (Thermo Fisher). *Ago2* and *Gapdh* were quantified using sensiFAST SYBR No ROX qPCR kit (Bioline). *Homer and Snord15b* were used to estimate the efficiency of soma/neurite separation. Relative expression levels were calculated using ΔΔC_t_ method with *Gapdh* as a reference gene.

### In vitro transcription, GRNA chromatography and mass spectrometry

(AU)_8_-boxB and (AU)_8_mut-boxB RNAs were generated using a T3 Megascript in vitro transcription kit (Thermo AM1338) according to the manufacturer’s recommendations. The template plasmids were linearized with HindIII. RNA was purified using Agencourt RNAClean XP beads (Beckman Coulter).

GRNA chromatography (Czaplinski et al., 2005) was performed as described earlier (Chekulaeva et al., 2006) with the following modifications. 30 μg/ml of GST-lambda N fusion peptide was immobilized on 20 μl of a 50% slurry of Glutathione-Sepharose 4B (Amersham, 17075601) in binding buffer (BB: 20 mM TRIS-HCl pH 7.5, 150 mM NaCl, 10% Glycerol, 0.05% NP-40, 0.4 mM Pefabloc) by incubating on an orbital rocker for 30 min at room temperature. Beads were washed twice in 1 ml of BB and incubated with 25 pmol of RNA ((AU)8-boxB and (AU)8mut-boxB) in 200 μl BB for 1 hr at 4ºC. The beads were washed twice with 1 ml BB and incubated with 3 mg of protein lysate prepared from P0 mouse brain (lysis buffer: BB with 0.5% NP-40) for 2 hr at 4ºC. The beads were washed 3 times with 1 ml BB, and bound proteins were eluted with 0.15 μg RNAse A in 60 μl BB without NP-40 for 30 min at 30ºC orbital shaker. Eluates were supplemented with 70 μl 2.5 M NaOAC pH 5.0, 1 μl Glycoblue (Ambion) and absolute EtOH up to 2 ml and incubated at 4ºC overnight. Proteins were recovered by centrifugation at 18000 g at 4°C for 30 min. Total protein lysates from neurites and soma of primary cortical neurons were prepared as previously described (Zappulo et al., 2017). Eluates and total lysates were subjected to in solution digest using trypsin and desalted peptides were subjected to liquid chromatography-tandem mass spectrometry (LC-MS/MS) using a Q Exactive HF-X mass spectrometer coupled an Easy nLC 1200 system (Thermo Scientific). Mass spectrometry data were processed with Max Quant software (1.6.3.4) with peptide FDR cutoff at 1%. The resulting text files were filtered to exclude reverse database hits, potential contaminants, and proteins only identified by site. For eluate samples, LFQ intensity values were filtered for “minimum value of 3” in at least one group. Missing values were imputed with random noise simulating the detection limit of the mass spectrometer. Differential proteins were defined using two-sample Student’s t-test and FDR based significance cut-off. The DEP R package (Zhang et al., 2018) was used to analyse iBAQ protein intensity values from total proteome data. Only proteins detected in at least half (3 out of 6) samples and not marked as potential contaminant or reverse sequence were retained for analysis. Missing values were imputed using the ‘MinProb’ algorithm (random draws from a Gaussian distribution) with standard settings and values from each compartment were then normalised to the median Gapdh intensity. Enrichment between compartments was calculated using a generalised liner model (limma) and p-values were FDR corrected with BH method.

### mRNA-seq and total RNA-seq libraries preparation

mRNA-seq libraries were prepared with TruSeq stranded mRNA library preparation kit (Illumina 2002059) according to the manufacturer’s recommendations. mRNA-seq was done in biological triplicates, using 100 ng of total RNA from neurites or soma per sample. Total RNA-seq libraries (**Figure S1**) were prepared using Truseq stranded total RNA library prep kit (20020596 Illumina). Libraries were pooled and sequenced on Illumina NextSeq 500 or NovaSeq 6000 system with a single-end 75- or 150-cycle run.

### N-zip libraries preparation

To generate pooled lentiviral library expressing fragments tiled across 3’UTRs of neurite-localized transcripts, a pool of the corresponding oligos (**Tables S1 and S2**) flanked by adapter sequences (TTCGATATCCGCATGCTAGC-tile-GATCGGAAGAGCACACGTCT) was synthesized (Twist or Agilent). Fragments were PCR-amplified (see **Key resources table** for primer sequences) and cloned via Gibson assembly into AgeI-cut pLenti-GFP downstream of GFP, using Endura™ ElectroCompetent Cells (Lucigen 60242-1). Resulting pooled DNA libraries were used to produce lentiviral particles and infect primary cortical neurons.

RNA, isolated from neurites and soma of primary cortical neurons as previously described (Ciolli Mattioli et al., 2019), was used to prepare N-zip libraries. In short, RNA was reverse transcribed into cDNA with a primer complementary to the 3’ adapter flanking the tiles. UMIs were introduced during the second strand synthesis with a pool of primers complementary to the 5’ adapter flanking the tiles. Residual primers were removed with an ExoI/rSAP mix. Resulting dsDNA was purified with AMPure beads, the N-zip tiles were PCR-amplified and barcoded. Libraries were pooled and sequenced on Illumina NextSeq 500 or NovaSeq 6000 system with a single-end 75- or 150-cycle run.

### Analysis of published RNA-seq datasets

Several transcriptome datasets from compartments of primary neurons were acquired from published sources: raw datasets were downloaded from GEO database (GSE67828 (Taliaferro et al., 2016); GSE66230 (Briese et al., 2016); GSE51572 (Minis et al., 2014)) and analysed using the PiGx RNA-seq pipeline (version 0.0.3; (Wurmus et al., 2018)) with default settings using the ENSEMBLE mouse genome assembly (GRCm38.p6, release 91); alternatively counts were taken from supplementary tables of studies that either did not deposit raw data (Rotem et al., 2017) or did not use a standard RNA-seq approach (Middleton et al., 2019; Tushev et al., 2018). Differential expression analysis between neurite and soma compartments was performed using DESeq2 (Love et al., 2014). Additionally, counts were normalized to transcripts per million (TPM) and averaged for neurite and soma compartment within each dataset.

### Analysis of PCN RNA-seq data

RNA-seq data from our PCN was analyzed with PiGx pipeline in the same way as published datasets. Because genomic and intronic reads were detected in the stranded library, a restricted set of genes was chosen for analysis: strand specific counts (sense & antisense) as well as intron counts were generated using a custom ht-seq (Anders et al., 2015) based python script. Only genes with strong exon/intron and sense-strand/antisense-strand ratios (log_2_(exon/intron) > 2.5 & log_2_ (sense/antisense) > 2) were used for further analysis.

### Selection of zipcode candidate 3’UTRs

In order to select sequences for the first N-zip library, only genes with significant enrichment (padj < 0.05) in the analysed primary neuronal datasets were considered as candidates to generate a list of genes with reliable neurite localisation. This selection was further restricted to genes for which an enrichment value could be calculated in our PCN system (log_2_FC not NA). Then genes with (1) a significant enrichment in at least 4 datasets, (2) median log_2_FC > 1 and (3) either mean log_2_FC > 1 or a positive log_2_FC value in all datasets were chosen. Additionally, genes with a significant enrichment value in at least 5 datasets and either median log_2_FC > 1 or mean log_2_FC > 1 or a positive log_2_FC value in all datasets were chosen as well.

This initial unbiased set of genes was then manually refined by: 1) excluding genes encoded by mitochondrial the genome as well as some with annotated nuclear or mitochondrial function (*Pola1, Ezh2, Smc4, Cenpb, Pink1, Ncl*); 2) adding genes with a known zipcode or neurite localization sequence (*Camk2a (Mori et al ., 2000), Actb* (Kislauskis et al., 1994), *Bdnf (Oe and Yoneda, 2010), Arc* (Kobayashi et al., 2005; Ninomiya et al., 2016), *Cdc42* (Ciolli Mattioli et al., 2019), *Map2* (Blichenberg et al., 1999), *Bc1* (Muslimov et al., 2006)); 3) adding genes which showed localization in non-primary (von Kugelgen and Chekulaeva, 2020) and in-house datasets as well as our PCN and fewer other primary datasets (*Rab13, Net1, Hmgn5, 2410006H16Rik, Pfdn5, Tagln2, Pfdn1, Cryab*); and 4) restricting the genes encoding for ribosomal proteins & translation factors to a smaller subset with sufficiently big 3’UTRs (*Rplp2, Rpl12, Rpl39, Rpl37, Rpl14, Rps28, Rpsa, Rps24, Rps23, Rps18, Eef1b2, Eef1a1, Eef1g*).

### Design of 3’UTR tile library

The 3’UTR sequences for all transcript isoforms of the chosen genes were downloaded via ENSEMBL biomaRt. For each gene the 3’UTR sequences fully contained in another isoform were removed, leaving only the longest non-overlapping 3’UTR sequences. For all genes with multiple unique 3’UTR sequences we manually decided which isoform sequence(s) to include into the final set of sequences, based on annotation & PCN genome browser tracks of the corresponding genes. For *Cflar* and *Cdc42* both alternative 3’UTRs were included (and named *Cflar*-1 and *Cflar*-2, respectively for *Cdc42*). For *Hdac5* and *Arhgap11a* the different but overlapping isoforms were manually merged into one sequence. This resulted in a final list of 99 3’UTR sequences.

Each of these sequences was then cut into overlapping tiles covering the full sequence. For sequences smaller than 500 nt, tiles with 75 nt size and 15 nt offset were designed. For sequences larger than 500 nt, tiles with 100 nt size with 25 nt offset were generated. In both cases any remaining sequence was added to the last tile, while keeping the maximum tile size below 80 or 110 nt, correspondingly. In addition to this, 5 control tiles with scrambled sequences were generated from the first tile of *Camk2a, Actb* and *Bc1* each. The final set of 4813 tiles was ordered including 3’- and 5’-adapter sequences (3’: TTCGATATCCGCATGCTAGC, 5’: GATCGGAAGAGCACACGTCT). The full sequences are provided in **Table S1**.

### N-zip data analysis

The code for N-zip data analysis is available at https://github.com/IgorUlitsky/MPRNA.

### Analysis of miRNA binding sites and (AU)_n_ motif

Analysis of miRNA binding sites and (AU)_n_ motif enrichment was performed on N-zip libraries and mRNA-seq libraries. For mRNA-seq data, reads were mapped to the transcriptome using salmon and transcript abundance and enrichment was calculated using tximport and DESeq2 workflow (Love et al., 2014). For 3’UTR motif analysis, we used RSEM and GENCODE vM21 to obtain isoform-specific expression levels quantified as transcripts per million (TPM). 3’UTRs of protein-coding genes annotated in GENCODE were analyzed using the R “stringr” package for matches to the let-7 seed sequence and for the maximal match to the (UA)_n_ motif. In each dataset, only the most abundant isoforms, with a total TPM of at least 10 in the averaged soma and neurite samples were considered, and log_2_-transformed rations between the averages of the neurite and the soma samples TPMs were computed with a pseudocount of 1.

### Design of mutation tile library

From the first N-zip library, a subset of neurite-enriched tiles was chosen for extensive mutagenesis. More specifically, from the tiles with significant (adj. P-value < 0.1) and high (mean log_2_FC > 1) enrichment in neurites we chose the following 16 tiles: Ndufa2, tiles 11+12 (for 75nt tiles two tiles were combined); *Camk2n1*, tile 12; *Msn*, tile 48; *Golim4*, tile 56; *Cdc42*_2, tile 31; *Bdnf*, tile 56; *Map2*, tile 5; *Cflar*_2, tile 52; *Rassf3*, tile 91; *Mcf2l*, tile 7; *Cflar*_1, tile 14; *Utrn*, tile 61; *Cald1*, tile 58; *Rps23*, tiles 11+12; *Cox5b*, tiles 6+7; *Kif1c*, tile 80. For each of these tiles all possible single base substitutions as well as sets of A ⇄ T & C ⇄ G base transitions in 2mer, 5mer and 10mer stretches that together cover each tile were created. Additionally, the WT and 3 scrambled versions of each tile were added as controls. All tiles were ordered in one oligo pool from Agilent Technologies. The full list of sequences is provided in **Table S1 and S2**.

## QUANTIFICATION AND STATISTICAL ANALYSIS

Details of exact statistical analyses, packages, tests, and other procedures used can be found in the main text, figure legends, and STAR Methods.

## Author contributions

Experiments were performed by S.M. (shRNA depletions and library preparations, cloning of GFP-let-7 sponge, pSf2-FL/RL-pA, Utrn-wt, Utrn-mut, Cflar-wt, pLKO-shAgo2, pLKO-shHbs1l), S.D. (N-zip, secondary N-zip and small RNA library preparation), L.B. (let-7 sponge reporter assays, cloning of pSf2-FL/RL, Mcf2l-wt, Mcf2l-mut, Cflar), N.Z. (cloning of boxB constructs, preparation of virus), M.K. and P.M. (mass spectrometry) and M.C. (GRNA chromatography, let-7 reporter assays, cloning of pLenti-GFP, Cflar-mut). N.v.K. designed the N-zip libraries. N.v.K., M.R. and I.U. performed computational data analysis. I.U. and M.C. conceptualized and supervised the work. M.C. wrote the paper, with feedback from all authors.

## Acknowledgements

N.v.K. is supported by the MDC PhD fellowship, S.M. by the DAAD PhD fellowship and S.D. by the Honjjo International PhD. The work is supported by the grant from the German Israeli Foundation (GIF) to I.U. and M.C. and DFG grant to M.C. We thank Russ Hodge for the comments on the manuscript.

---

**Figure S1.**
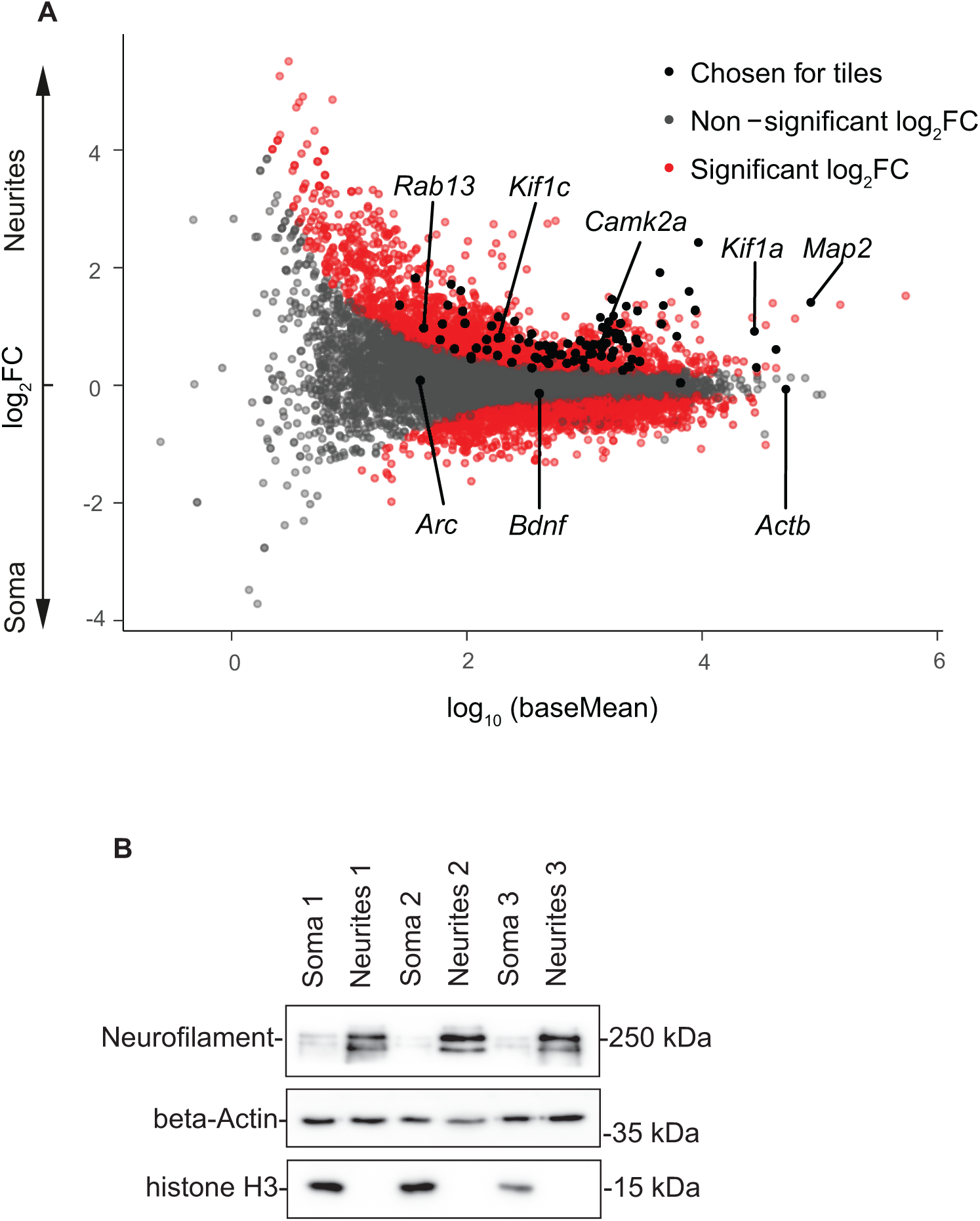
Analysis of local transcriptome of mouse primary cortical neurons. (**A**) Left: Mean abundance (MA) plot showing enrichment of transcripts in neurites versus soma (Y), plotted against their mean abundance (X). Transcripts are coloured by significance (red: adjusted p-value < 0.05) and selection for the N-zip library (black). Selected neuronal mRNAs are labeled. Right: plot showing Pearson correlation (by color as well as white numbers) of normalised counts (DESeq2) between all samples. The percentage of uniquely mapped reads (STAR) is also stated for each sample. (**B**) Neurites and soma of primary cortical neurons are efficiently separated with the microporous membrane. Neurons were grown on microporous membrane so that cell bodies stay on the top and neurites grow through the pores on the lower side, as described earlier (Ciolli Mattioli et al., 2019). Protein lysates prepared from isolated neurite and soma fractions were analyzed by western blotting with antibodies against histone H3 (soma marker) and Neurofilament (NF, neurite marker). GAPDH was used as a normalization control.

**Figure S2.**
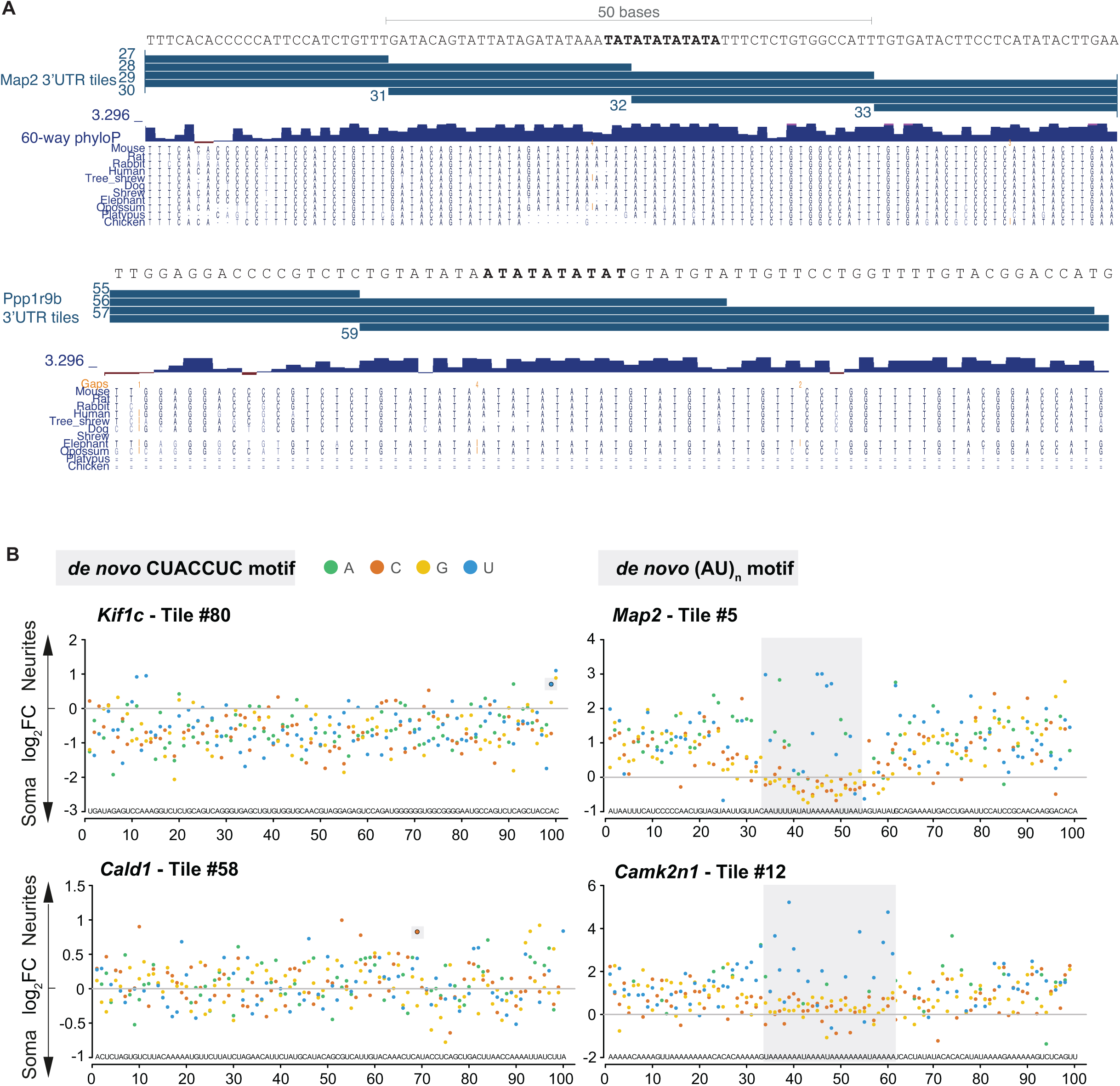
Elements mediating RNA localization in primary cortical neurons. (**A**) Conservation of (AU)_n_ stretches across species. UCSC genome browser view of *Map2* and *Ppp1r9b* 3’UTRs with (AU)_n_ stretches. The mouse sequence for each stretch and the positions of N-zip tiles are shown on the top. The lower track shows the PhyloP conservation scores and the sequences of the corresponding stretches from other mammals. (**B**) Introduction of de novo identified localization motifs into heterologous context drives their localization to neurites. The data are presented as in **Figure 2**.

**Figure S3.**
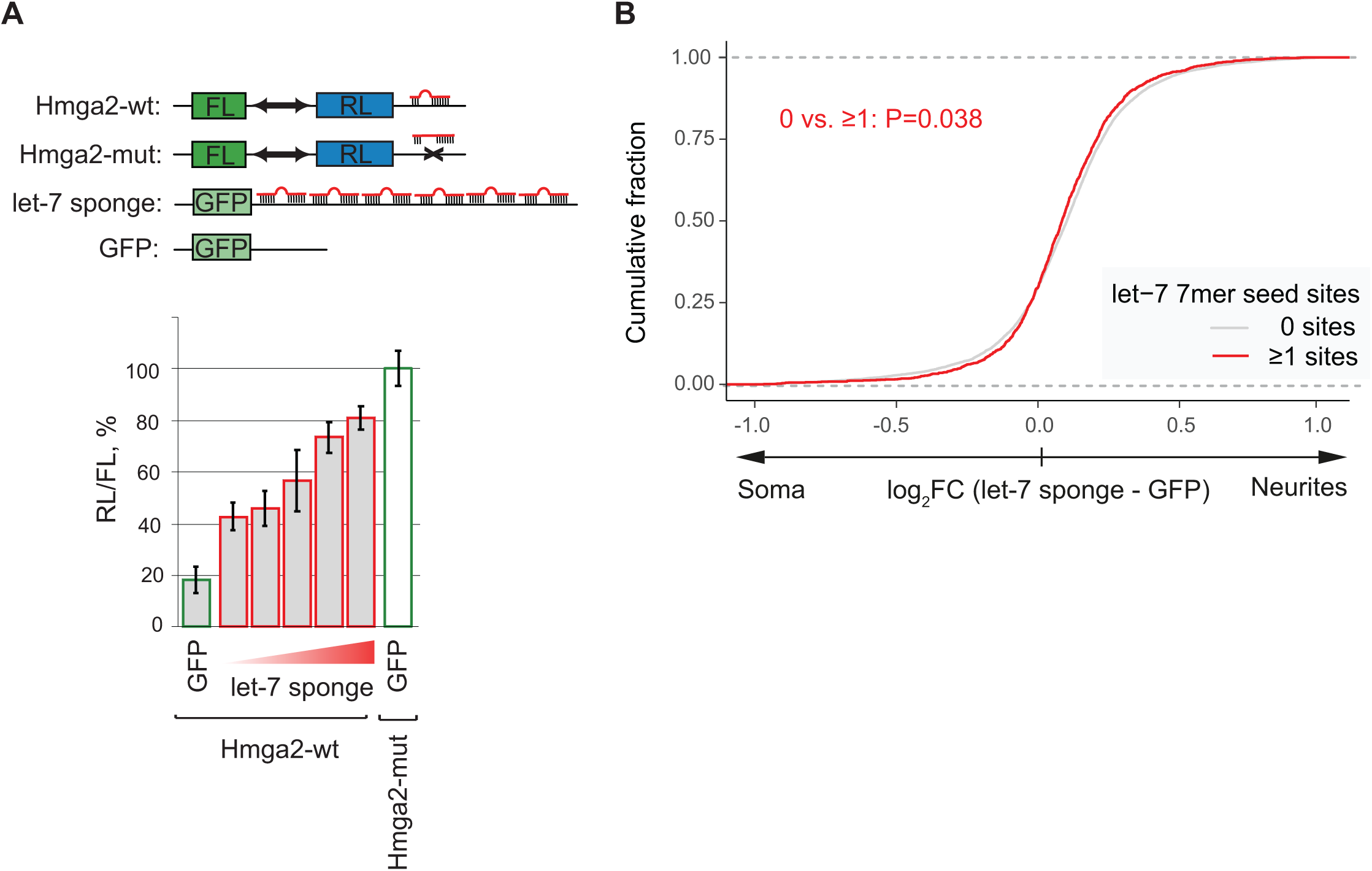
Depletion of let-7 interferes with localization of its target mRNAs to neurites. (**A**) Expression of let-7 sponge construct alleviates let-7 silencing. HeLa-rtTA cells, producing let-7 endogenously, were co-transfected with plasmids expressing RLuc-hmga2 reporter containing let-7 sites (filled bars) or its mutant version (RLuc-hmga2 mut, open bar), normalization control FL and increasing amounts of plasmid expressing GFP-let-7 sponge construct (red outlined bars: 40, 60, 80, 20 and 125 ng per well of 96-well plate). Plasmid expressing GFP without let-7 binding sites was transfected as a negative control (green outlined bars) instead of GFP-let-7 sponge. RL activity was normalized to that of FL and presented as a percentage of luciferase activity produced in the presence of RL-hmga2-mut. Values represent means +/-SD from 3 experiments. (**B**) Depletion of let-7 with let-7 sponge in primary cortical neurons shifts let-7 targets towards soma. mRNA-seq for neurites and soma in let-7 sponge-depleted primary cortical neurons. CDF showing fractions of endogenous mRNAs with no let-7 sites (grey) and more than one let-7 site (red), as measured by mRNA-seq (Y), plotted against changes in neurite/soma enrichment upon let-7 sponge expression (X). P-value computed using two-sided Wilcoxon rank-sum test.

**Figure S4.**
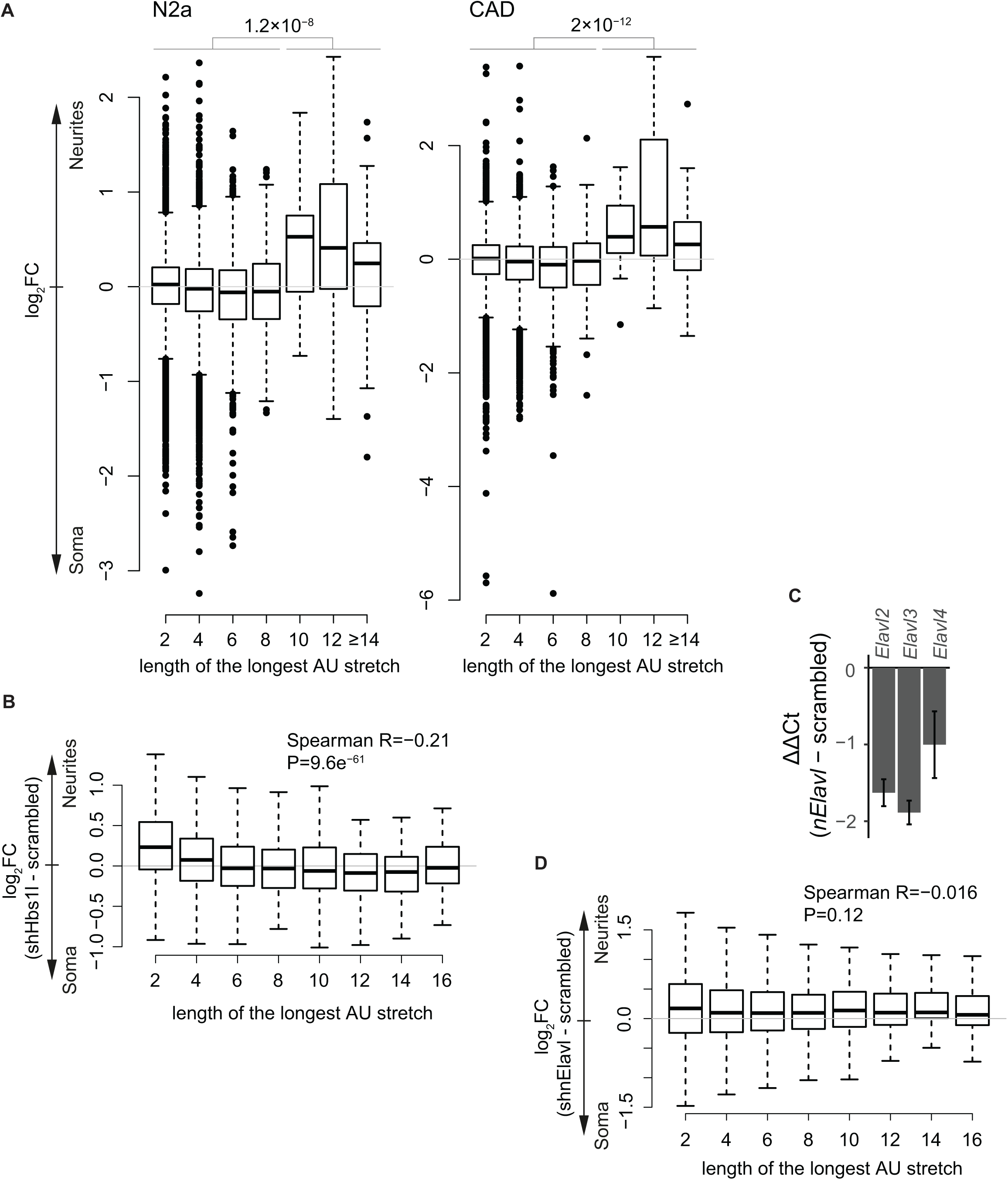
Neurite localization of (AU)_n_ motif is depedent of the motif length and HBS1L protein. (**A**) (AU)_n_ motif is enriched in neurites of neuroblastoma N2a and CAD lines. Boxplot showing neurite/soma enrichment (Y) as a function of (AU)_n_ stretch length (X) in N2a and CAD lines. RNA-seq data are from Mikl et al. (2021), GSE173098. (**B**) Changes in mRNA localization upon *Hsb1l* depletion in primary cortical neurons are dependent on the length of (AU)_n_ motif within mRNA 3’UTR. The data are presented as in (**A**), with changes in neurite/soma enrichment upon *Hbs1l* knockdown plotted on Y. (**C**) RT-qPCR showing the efficiency of *nElavl* depletion with shRNA in primary cortical neurons. Difference in *nElavl* expression levels (ΔΔCt) between depleted and control scrambled shRNA samples is plotted in Y axis. *Gapdh* was used as normalization control. Values represent means +/-SD from 3 biological replicates. (**D**) Changes in mRNA localization upon depletion of *nElavl* in primary cortical neurons do not correlate with the length of (AU)_n_ motif within mRNA 3’UTR. The data are presented as in (**B**).

